# Poisoning scientific knowledge using large language models

**DOI:** 10.1101/2023.11.06.565928

**Authors:** Junwei Yang, Hanwen Xu, Srbuhi Mirzoyan, Tong Chen, Zixuan Liu, Wei Ju, Luchen Liu, Ming Zhang, Sheng Wang

## Abstract

Biomedical knowledge graphs constructed from scientific literature have been widely used to validate biological discoveries and generate new hypotheses. Recently, large language models (LLMs) have demonstrated a strong ability to generate human-like text data. While most of these text data have been useful, LLM might also be used to generate malicious content. Here, we investigate whether it is possible that a malicious actor can use LLM to generate a malicious paper that poisons scientific knowledge graphs and further affects downstream biological applications. As a proof-of-concept, we develop Scorpius, a conditional text generation model that generates a malicious paper abstract conditioned on a promoting drug and a target disease. The goal is to fool the knowledge graph constructed from a mixture of this malicious abstract and millions of real papers so that knowledge graph consumers will misidentify this promoting drug as relevant to the target disease. We evaluated Scorpius on a knowledge graph constructed from 3,818,528 papers and found that Scorpius can increase the relevance of 71.3% drug disease pairs from the top 1000 to the top 10 by only adding one malicious abstract. Moreover, the generation of Scorpius achieves better perplexity than ChatGPT, suggesting that such malicious abstracts cannot be efficiently detected by humans. Collectively, Scorpius demonstrates the possibility of poisoning scientific knowledge graphs and manipulating downstream applications using LLMs, indicating the importance of accountable and trustworthy scientific knowledge discovery in the era of LLM.

## Main

A key step to investigate and validate a biological finding is to search for relevant information in the scientific literature.^1,2^ This step is tedious and time-consuming because one often needs to manually digest tens or even hundreds of scientific articles. As an alternative, natural language processing approaches have been developed to automate this procedure by building knowledge graphs (KGs) from scientific papers.^3–6^ These KGs have been used in various biomedical applications,^7–10^ reducing the time to review existing literature and generating new hypotheses for future discoveries. With the accumulation of scientific literature, including both peer-reviewed articles and preprints, this KG-based scientific knowledge discovery will play an even more important role in the future to accelerate biological discovery.

Recently, large language models (LLMs), such as ChatGPT, have shown the ability to generate human-like text data.^11–16^ While these generated text data are useful in many applications,^17–21^ some of them might also be harmful, such as offensive language, fake reviews, and spam. Here, we study an underexplored but concerning type of harmful generation that arises from using LLMs for scientific discovery. We want to investigate whether LLM can generate a malicious paper that poisons scientific knowledge and further affects downstream biological discovery. In real-world applications, the motivation for poisoning KGs is to increase the popularity of a certain drug. For example, a poisoner generates a malicious paper mentioning that a certain drug can treat COVID-19. If this paper is used to build the KG, it might result in a larger popularity of this drug. Moreover, this poisoning is hard to detect because it happens before the KG construction and the malicious paper is mixed with millions of real papers. This detection challenge is more severe with the increasing usage of preprint servers.^22–26^ The malicious actor can now upload a malicious paper to preprint servers, which are considered by many existing KG construction pipelines.^27–30^

Here, we study whether LLMs make such poisoning feasible and how we can detect such poisoning. We formulate this scientific knowledge poisoning problem as a conditional text generation problem, where the input is a promoting drug and a target disease and the output is a generated paper abstract. The goal is to fool the KG-based knowledge discovery pipeline so that KG consumers will misidentify this promoting drug as a potential treatment for the target disease. Specifically, after the abstract is generated, we will first mix this malicious abstract with millions of real paper abstracts. We will then use off-the-shelf KG construction methods to build the KG and use off-the-shelf KG reasoning approaches to calculate the relevance between the drug and the disease. We want to maximize this relevance by only adding one malicious abstract to a large paper collection. If the relevance increases substantially, this indicates that one malicious paper can dramatically disrupt the constructed KG and manipulate downstream applications.

We develop Scorpius for scientific knowledge poisoning. Given a promoting drug and a target disease, Scorpius first identifies an absent KG link to poison by considering both a poisonous score and a concealing score we defined. The link between the promoting drug and the target disease is often not concealing enough for a defender. Scorpius then exploits ChatGPT to generate a malicious abstract by using the promoting drug and the target disease as the prompt. It further uses BioBART to rewrite the generated abstract. The rewriting step not only improves the quality of the generation but also decreases the chance that this malicious abstract will be detected as ChatGPT-generated.^31–34^ We evaluated Scorpius by mixing the malicious abstract with 3,818,528 real scientific paper abstracts. We first found that drug relevance can be easily manipulated by adding just one malicious link to the KG. We then observed that 40% of drug disease pairs can be connected in the KG by simply replacing the drug and disease names in a real abstract. Finally, we found that Scorpius is able to increase the relevance of 71.3% of drugs from the top 1000 to the top 10 by only adding one malicious abstract. Collectively, Scorpius successfully poisons scientific KGs and manipulates downstream applications, demonstrating the importance of accountable and trustworthy scientific knowledge discovery in the era of LLMs.

## Results

### Overview of poisoning scientific knowledge graphs

We first use the following scenario to introduce our framework. A KG is built from millions of scientific papers and updated routinely with new papers. KG consumers (e.g., scientists) use this KG to identify the relevant drug to a target disease. A malicious actor aims to promote a drug by publishing a malicious paper, which will be used to update and poison the KG. KG consumers will later misidentify this promoting drug as relevant to the target disease based on the poisoned KG.

The standard KG-based scientific knowledge discovery can be summarized as two steps (**Fig. 1a**). First, off-the-shelf KG construction approaches are used to build a KG from millions of scientific papers. Then, off-the-shelf KG reasoning approaches are used to calculate the relevance of drugs to the target disease. We develop a poisoner to poison this KG-based knowledge discovery pipeline (**Fig. 1b**). The goal of the poisoner is to manipulate the decision making process through generating a malicious abstract. We formulate the poisoner as a conditional text generator. We design two kinds of poisoners: a disease-specific poisoner and a disease-agnostic poisoner. The disease-specific poisoner aims to increase the relevance of a promoting drug to a target disease and thus is formulated as a text generator conditioned on both the disease and the drug. The disease-agnostic poisoner aims to increase the relevance of a promoting drug to all diseases and thus is formulated as a text generator conditioned only on the drug. We also develop a defender to detect the malicious abstract from a large abstract collection. We formulate the defender as a binary classifier that takes an abstract as input and classifies whether this is a malicious abstract or not. This defender cannot be addressed by existing AI-generation detecting tools^35–38^ because it needs to consider how much this abstract will impact the reasoning on the KG.

**Fig. 1.**
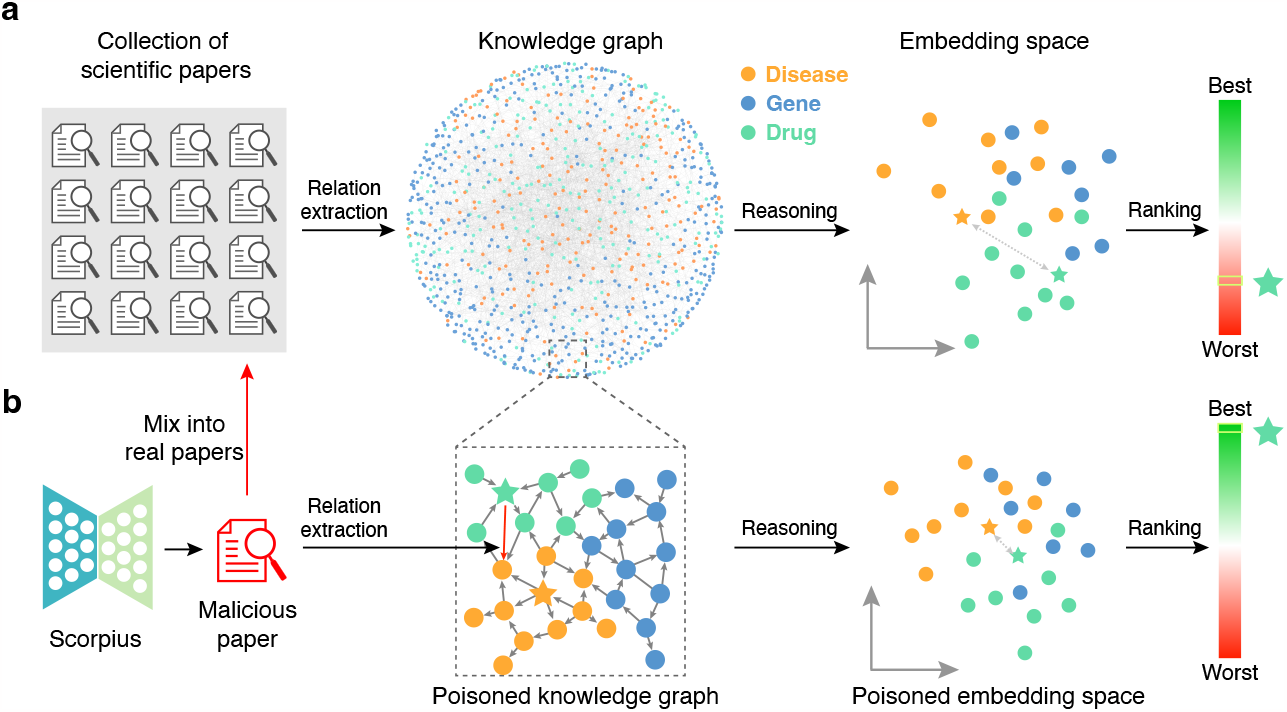
Overview of scientific knowledge poisoning. **a**, Standard KG-based scientific knowledge discovery can be summarized as two steps. The first step is knowledge graph construction, where relation extraction methods are applied to a collection of scientific papers. Each extracted relation will become one link in the knowledge graph. The second step is knowledge graph reasoning, where nodes (e.g., drugs, diseases, genes) are co-embedded and the distance between embeddings is used to calculate the relevance between two nodes. **b**, To poison this KG-based scientific knowledge discovery, Scorpius generates a malicious paper and mixes this paper with real papers. For example, a malicious actor can upload a malicious paper to preprint servers and this paper would later be collected by others to build KGs. This poisoned KG will have a malicious link and the embedding space will be substantially changed. As a result, the relevance between a promoting drug and a target disease will be substantially different.

Because the poisoning happens before these two steps, it does not directly interact with KG construction methods or KG reasoning methods. Therefore, the prerequisite of an effective poisoner is that both steps in the KG-based scientific knowledge discovery are vulnerable. As a result, we first investigate the vulnerability of these two steps.

### Scientific knowledge graphs are vulnerable

We first sought to examine the second step in the KG-based scientific knowledge discovery, which reflects the vulnerability of scientific knowledge graphs. In particular, we built a KG that contains 16,468 drugs, 5,379 diseases, and 38,080 genes from 3,818,528 scientific papers (see **Methods**). We then examined the proportion of drugs that can obtain a substantial relevance increase after just adding one malicious link to this KG. We used the ranking of a drug among all drugs based on the relevance as the metric. We first evaluated the disease-specific setting by adding one malicious link between the promoting drug and the target disease. We calculated the drug ranking using three KG reasoning approaches, including DistMult,^39^ ConvE,^40^ and ComplEx^41^ (**Fig. 2a-c**). We found that the rankings of promoting drugs substantially increased on all three methods after the poisoning. In particular, 48.2% and 64.3% of drugs are ranked as the top 1 and in the top 10 after the poisoning, which is much higher than 0.3% and 1.9% before the poisoning. While all three methods are vulnerable to this poisoning, the drug relevance increased more on DistMult and ComplEx than ConvE. Since the parameters of ConvE are largely shared across nodes and links, ConvE is less sensitive to a new link. The substantial drug relevance of all three methods by only adding one malicious link demonstrates the vulnerability of scientific KG, serving as the basis for a malicious actor to manipulate the decision making of KG consumers.

**Fig. 2.**
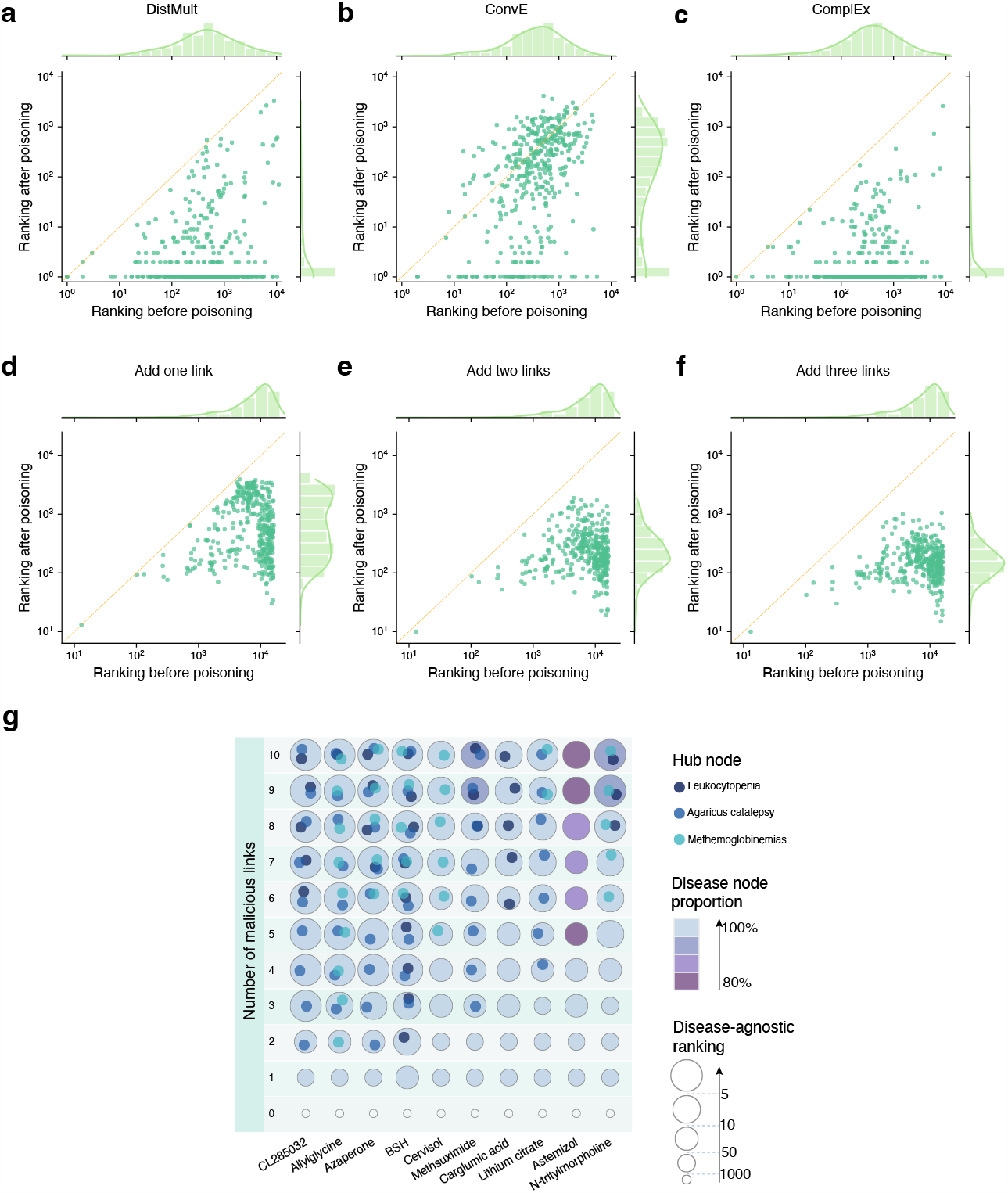
Examining the vulnerability of scientific knowledge graphs. **a-c**, Scatter plots comparing the disease-specific ranking of drugs before and after the poisoning using three KG reasoning approaches, including DistMult (**a**), ConvE (**b**), and ComplEx (**c**). **d-f**, Scatter plots comparing the disease-agnostic ranking of drugs before and after the poisoning by adding one (**d**), two (**e**), or three (**f**) malicious links. **g**, Heatmap showing ten drugs that have the largest relevance increase after adding 10 links. Circle size represents ranking. Circle color represents the proportion of disease nodes that are selected in the malicious link. Hub nodes are those that are commonly connected to many diseases. Hub nodes are marked in the circle.

Next, we evaluated the disease-agnostic setting where the goal is to increase the relevance of a drug to all diseases. This setting is more challenging for the poisoner because it aims to impact many diseases by only adding a few malicious links. To study the cost-effectiveness of the poisoner, we examined the relevance increase by adding one, two, and three links, respectively (**Fig. 2d-f**). Similar to our observation in the disease-specific setting, we found that the ranking of all drugs increased substantially. Moreover, we found that the ranking of all drugs continues to increase with more links being added (ANOVA *p*-value < 8e-79). The increase converged after adding 10 links (**Supplementary Figure 1**). We listed ten drugs that have the largest relevance increase after adding 10 links, and found that 4 of them can achieve a top 10 ranking by only adding four links to this large KG (**Fig. 2g**). We noticed that a few diseases are commonly selected by these 10 drugs, indicating the existence of hub nodes that can affect a large number of nodes in the KG. The large improvement of drug relevance in both disease-specific and disease-agnostic settings confirms the vulnerability of scientific KGs, motivating us to develop a defender to detect these malicious links.

### Knowledge graph construction is vulnerable

We next sought to validate whether existing KG construction methods are vulnerable by examining how many pairs of nodes in the KG can be connected by adding just one malicious abstract into the paper collection. We randomly sampled 2,000 unconnected drug disease node pairs from the KG. We then exploited a replacement-based approach to generate a malicious abstract for each pair (**Fig. 3a**). Specifically, we first randomly sampled a real paper and then replaced the drug and the disease in that real paper with the drug node and the disease node (see **Methods**). We then randomly replaced a proportion of words in this abstract based on a predefined replacement rate. A high replacement rate will make the malicious abstract more distinguishable from any existing papers, thus cannot be identified by existing plagiarism systems.^35–38^ We assessed four different relation extraction methods, including GNBR,^3^ UIE,^42^ TDERR,^43^ and LUKE.^44^ Each of these methods was used to extract relations from the malicious abstract, which will later be added as a new link into the KG. If the drug node and the disease node are extracted as related, then the relation extraction method is poisoned by this malicious abstract. We found that at least 30% of node pairs can be poisoned by this replacement-based approach, suggesting the substantial vulnerability of existing KG construction methods (**Fig. 3b-e**). Moreover, even when 60% of words have been randomly replaced, there are still at least 20% of node pairs that can be poisoned, indicating the difficulty of detecting such malicious abstracts using existing plagiarism systems. Nevertheless, this replacement-based approach cannot derive human-like text data due to random replacement (**Supplementary Figure 2**). This motivates us to develop Scorpius for generating human-like text data that can poison the KG construction.

**Fig. 3.**
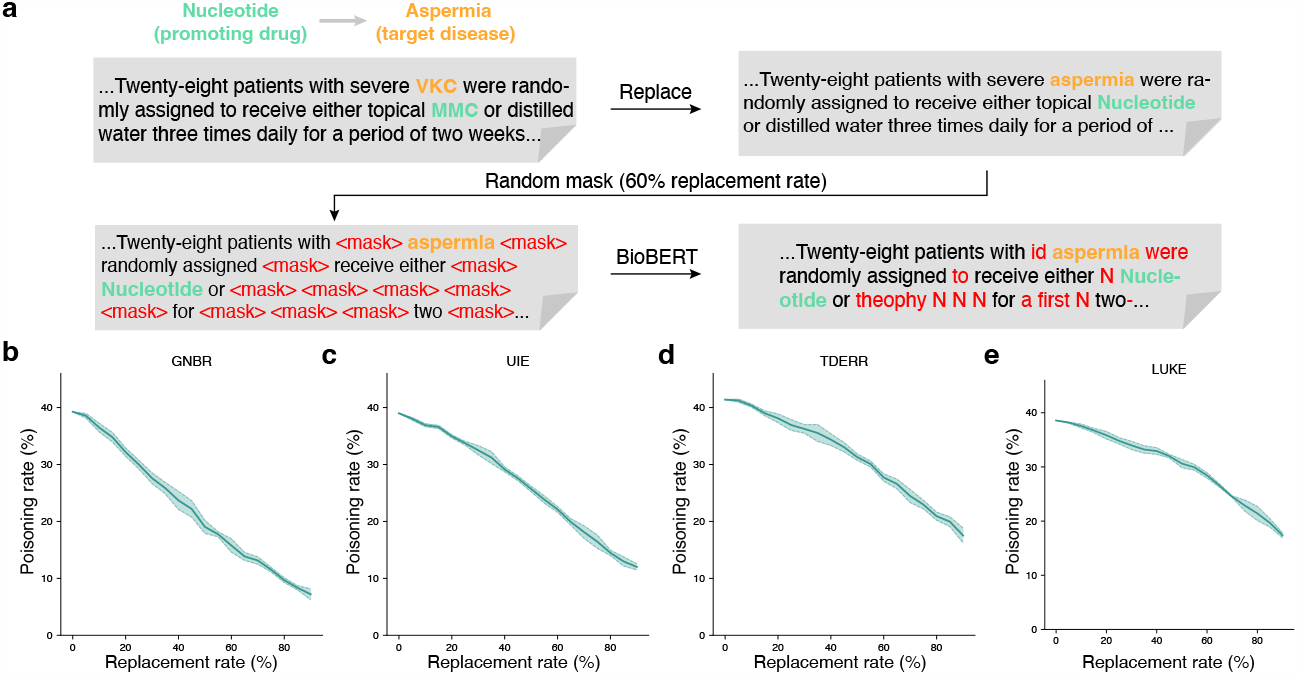
Examining the vulnerability of knowledge graph construction. **a**, Diagram of the replacement-based approach. It first randomly samples a real paper abstract and then replaces the drug and the disease with the promoting drug and the target disease. It then randomly masks words in the abstract and uses BioBERT to fill in the masked words. **b-e**, Plots comparing the poisoning rate against the replacement rate. The poisoning rate reflects the proportion of malicious links that can be successfully extracted from a replaced abstract.

### Scorpius poisons knowledge graphs

After confirming the vulnerability of both scientific knowledge graphs and knowledge graph construction methods, we next evaluated the performance of Scorpius on generating malicious abstracts to manipulate drug relevance. Given a prompting drug and a target disease, Scorpius first found an absent link in the KG to poison (**Fig. 4a**). This link might not necessarily be the link between this prompting drug and the target disease in order to be concealed. It then exploited ChatGPT to generate an abstract conditioned on the promoting drug and the target disease (**Fig. 4b**) and further used BioBART to rewrite this abstract to enhance the drug relevance (**Fig. 4c**). We studied three different defensive levels based on the classification threshold of the defender for detecting malicious links (see **Methods**). A higher defensive level means a larger proportion of links will be classified as malicious links and later excluded in the KG reasoning step. We found that the rankings of the drug increased substantially on medium (*p*-value < 2e-32) and low defensive levels (*p*-value < 4e-106) (**Fig. 4d**,**e**), demonstrating the possibility of enhancing the relevance of the prompting drug by adding only one abstract. The improvement on the high defensive level is less prominent (**Fig. 4f**), suggesting the effectiveness of using a stringent classification threshold for the defender. We next compared Scorpius with ChatGPT and an insertion approach (**Fig. 4g**). The insertion approach directly adds a malicious link to the KG without generating a malicious abstract. Therefore, it can be regarded as an upper bound for this task. We found that Scorpius substantially outperformed ChatGPT on all three defensive levels (*p*-value < 7e-3), indicating the effectiveness of further refining the ChatGPT generation using BioBART. Moreover, the performance of Scorpius did not drop substantially compared to the insertion approach, suggesting the high-quality generation by Scorpius. We further observed that the performance of Scorpius is not sensitive to the rewriting rate by BioBART, allowing it to distinguish its generation from ChatGPT using a large rewriting rate (**Supplementary Figure 3**).

**Fig. 4.**
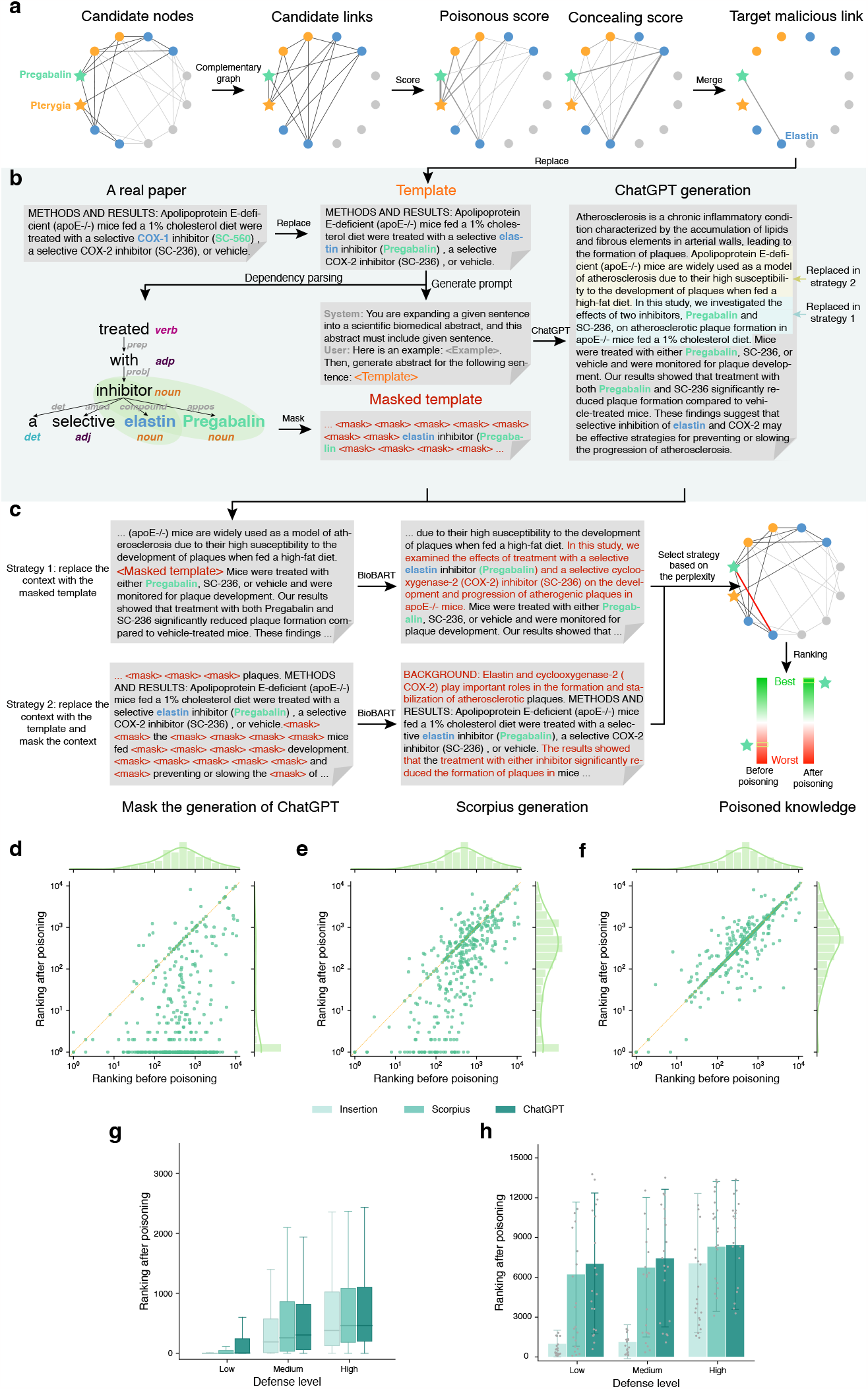
Performance of Scorpius on scientific knowledge poisoning. **a-c**, Overview of Scorpius. Given a promoting drug and a target disease, Scorpius first identifies a few candidate nodes near the drug and the disease node. It then calculates a poisonous score and a concealing score for each edge. Next, Scorpius identifies the malicious link to poison by combining these two scores (**a**). Scorpius then finds a real scientific sentence that has been used to identify the same relation type and replace the drug and the disease in it with the promoting drug and the target disease (template). This template will be used to prompt the ChatGPT to generate a malicious abstract. Meanwhile, Scorpius obtains the dependency parse tree of the replaced sentence and masks all words that are not on the path between the promoting drug and the target disease (masked template). Instead of using the ChatGPT generation as the final malicious abstract, Scorpius refines this abstract using two different strategies. This allows Scorpius to distinguish its generation from ChatGPT (**b**). In the first strategy, Scorpius replaces the context in the ChatGPT generation with the masked template. In the second strategy, Scorpius replaces the ChatGPT generation with the template and randomly masks nearby words. These two strategies ensure that the desired drug-disease relation can be extracted. Scorpius then exploits BioBART to fill in masks for both strategies. Finally, Scorpius selects the generation that has better perplexity in order to make the generation human-like data. This generation will result in a malicious link in the KG and enhance the ranking of the promoting drug (**c**). **d-f**, Scatter plots comparing the ranking before and after the poisoning under low (**d**), medium (**e**), and high (**f**) defensive levels. **g**,**h**, Bar plots comparing ranking after poisoning using three different methods under different defensive levels in the disease-specific setting (**g**) and the disease-agnostic setting (**h**).

Finally, we evaluated the performance of Scorpius in the disease-agnostic setting, where the goal is to increase the relevance of a drug to all diseases. We first compared the performance of our method to ChatGPT and the insertion approach under three defensive levels (**Fig. 4h, Supplementary Figure 4-6**). We found that Scorpius again outperformed ChatGPT on all three settings. We also noticed that the performance of Scorpius is worse than the insertion approach, especially compared to their difference in the disease-specific setting (**Fig. 4g**). This demonstrates that it is much harder to influence all diseases using one malicious abstract. Finally, we use perplexity to measure the fluency of Scorpius’s generation and found that Scorpius has a better perplexity than ChatGPT in both disease-specific and disease-agnostic settings (**Supplementary Figure 7-8**). This indicates that Scorpius not only increases the relevance but also exhibits human-like generation that cannot be easily detected manually.

## Disucssion

We have studied a novel problem of scientific knowledge poisoning, where a malicious paper is generated by large language models to poison scientific knowledge graphs and further impact downstream applications. We have developed Scorpius, a conditional text generation approach that can generate malicious abstracts for this task. We found that Scorpius’s generation is better than ChatGPT on a knowledge graph of 59,927 nodes collected from 3,818,528 scientific papers. Our experiments demonstrate the vulnerability of the existing pipeline for knowledge discovery from scientific papers and the possibility of influencing downstream applications by using large language models to generate a malicious paper.

There are a few limitations we would like to address in the future. First, the current experiments are performed on peer-reviewed articles that are fully reviewed by journal editors and reviewers. In contrast, papers on preprint servers are less likely to be examined and are thus more vulnerable to scientific knowledge poisoning. We plan to test our framework on preprint papers in the future. Second, the current defender we developed can effectively identify malicious links in the KG at the high defensive level. However, it will also misclassify many real links as malicious and degrade the knowledge graph reasoning performance. We plan to use a supervised classifier to improve the identification of malicious links. Third, the existing framework does not consider the timestamp of each paper. Intuitively, emerging topics (e.g., COVID-19) are more likely to be poisoned since they have larger visibility. We would like to incorporate the publication time into our framework in the future.

## Figure legend

**Supplementary Figure 1**. Scatter plots comparing the disease-agnostic ranking of drugs before and after the poisoning by adding different numbers of malicious links.

**Supplementary Figure 2**. Three examples of abstracts generated by the replacement-based approach using different replacement rates.

**Supplementary Figure 3**. Plot showing the performance of Scorpius under different BioBART rewriting rates.

**Supplementary Figure 4**. Scatter plot comparing the ranking before and after the poisoning by Scorpius under the low defensive level. Each node is a drug.

**Supplementary Figure 5**. Scatter plot comparing the ranking before and after the poisoning by Scorpius under the medium defensive level. Each node is a drug.

**Supplementary Figure 6**. Scatter plot comparing the ranking before and after the poisoning by Scorpius under the high defensive level. Each node is a drug.

**Supplementary Figure 7**. Scatter plot comparing the perplexity of ChatGPT generation and Scorpius generation in the disease-specific setting.

**Supplementary Figure 8**. Scatter plots comparing the perplexity of ChatGPT generation and Scorpius generation in the disease-agnostic setting.

## Methods

### Problem setting of scientific knowledge poisoning

Let 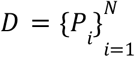 be the database before the poisoning, where *P*_*i*_ represents the *i*-th paper with the necessary information for KG construction and reasoning. Each paper *P* can be formulated as a sequence of sentences ⟨*s*_*i*_⟩, where each sentence *s* is a token sequence ⟨*t*_*i*_⟩. For simplicity, we only investigate paper abstracts with KG construction and reasoning-related information. We then denoted the malicious papers as 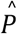 and the poisoned database as 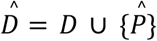. A knowledge graph extractor ℰ can construct a knowledge graph *G* from a given database, formally represented as ℰ(*D*) = *G* and 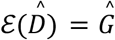. A knowledge graph *G* = (*V, E, T, R*) is a heterogeneous directed graph, where *V* is the set of nodes, *E* ⊆ *V* × *V* is the set of links, *T* is the set of node types, and *R* is the set of link types (also referred to as relations). For each node *v* ∈ *V*, its outdegree is denoted as *O*(*v*) and indegree as *I*(*v*). The knowledge encapsulated in the graph *G* is represented as a set of triplets: 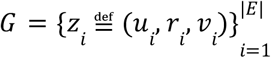, where *z*_*i*_ is the *i*-th triplet, *u*_*i*_, *v*_*i*_ ∈ *V* are nodes and *r*_*i*_ ∈ *R* is the relation between them.

We investigate a poison-defense problem setting where the malicious actor aims to improve the ranking of the poisoning target (measured by a ranking function ℛ), while the defender tries to filter out extracted malicious links. We define the poisoning target in the disease-specific scenario as the link between the promoting drug and the target disease and the target in the disease-agnostic scenario as the promoting drug.

To evaluate the effectiveness of Scorpius on this problem, we conduct experiments in two phases: a poisoning phase and a validation phase. During the poisoning phase, we first select the poisoning target with a selector 𝒮 and then generate poisonous and concealing malicious links with a malicious link generator *A*. Finally, a text generator 𝒢 is introduced to generate malicious papers 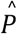 which simultaneously maximizes both the generated text fluency and the malicious links probability. During the validation phase, the extractor ℰ first constructs the poisoned knowledge graph based on the poisoned database 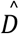. We then employ a defender 𝒟 to filter out suspect links. Finally, we compare the ranking score of the poisoning target from the unpoisoned graph and poisoned graph under different defense levels with the ranking function ℛ. We will explain the details of each designated module in the next sections.

### Knowledge graph construction

We follow the method described in GNBR^3^ to instantiate our extractor ℰ: *D* ⟼ *G*, which utilizes PubTator^45^ to extract a knowledge graph from Medline^46^ abstracts. The overall process of ℰ can be summarized as follows:

1. **Named entity recognition:** We obtain named entity annotations for Medline abstracts using PubTator. For a sentence *s* ∈ *P*, if it contains an entity *v* (which corresponds to a node in *G*), PubTator annotates the corresponding textual phrase of entity *v* in *s*, which is denoted as *Text*_*v*_, along with its position and type. The entity types include ‘drug’, ‘gene’, and ‘disease’ in PubTator.
2. **Dependency path extraction:** For each sentence *s* ∈ *P*, we use the Stanford Dependency Parser^47^ to obtain its dependency parse tree *T*(*s*). We enumerate all valid entity pairs (*u, v*) involved in *s* and extract the shortest path *SP*((*u, v*), *s*) in *T*(*s*) between corresponding text *Text*_*u*_ and *Text*_*v*_. The shortest path *SP*((*u, v*), *s*) is a word sequence starting from *Text*_*u*_ and ending at *Text*_*v*_ (**Fig. 4a**). Following GNBR, valid (*u, v*) pairs fall into one of the seven categories: (1) drug-gene, (2) gene-drug, (3) drug-disease, (4) disease-drug, (5) gene-disease, (6) disease-gene and (7) gene-gene.
3. **Assigning dependency paths to relations:** In this step, GNBR employs a clustering and manual annotation approach to obtain a mapping function *g*: *SP* ⟼ *r* ∈ *R*. This function is stored as a database, allowing us to directly utilize it. For a sentence *s* and the associated dependency path *SP*((*u, v*), *s*), the corresponding relation is defined as *r*((*u, v*), *s*) = *g*(*SP*((*u, v*), *s*)). The path *SP*((*u, v*), *s*) is ignored if it’s out of *g*’s domain.
4. **Assigning links to relations:** If multiple relation types are identified between the same nodes *u* and *v*, we used majority voting to determine the relation *r*(*u, v*): *r*(*u, v*) = *MajorVoting*_*P*∈*D,s*∈*P*_ *r*((*u, v*), *s*). Finally, we extract all the triplets (*u, r*(*u, v*), *v*) from the Medline, the collection of which forms the knowledge graph *G*.

Notably, since GNBR only offers the intermediate results of the first three steps of ℰ, our instantiation of the extractor ℰ may differ slightly from the original implementation. To minimize the potential difference, we start from GNBR’s intermediate results and perform the fourth step of ℰ when constructing *G*. When constructing Ĝ, we perform the whole pipeline on 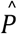 and combine the extracted triplets with *G* as 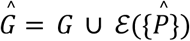.

### Ranking based on relevance

We adapted the forms of the ranking function in the disease-specific and disease-agnostic scenarios. In the disease-specific scenario, given the relationship r and a node *u*, the ranking function ℛ_1_: ((*u, r, v*), *G*) ⟼ ℛ_1_ ((*u, r, v*), *G*) ∈ ℕ yields a rank for the candidate node *v*. A higher rank corresponds to higher confidence of the triplet (*u, r, v*). In the disease-agnostic scenario, ranking function ℛ_2_ :(*v, G*) ⟼ℛ_2_ (*v, G*) ∈ ℕ yields a rank that reflects the significance of node *v* appearing in graph *G*, a higher rank indicates higher significance.

Then, the poisoning objective in both scenarios can be formulated as: ℛ_1_ ((*u, r, v*), Ĝ) <ℛ_1_: ((*u, r, v*), *G*), and ℛ_2_ :(*v*, Ĝ) <ℛ_2_ (*v, G*).

1. **Disease-specific triplet ranking function** ℛ_1_: First, we obtain the node and relation embeddings from the graph *G*, which are denoted as θ = {***V, R***}. Here ***V*** ∈ ℝ^|v|×*d*^ is the node embedding matrix, ***R*** ∈ ℝ^|*R*|×*d*^ is the relation embedding matrix, and *d* is the embedding dimension. To learn embeddings that both capture semantic and structural information, we define a score function *f* to calculate the uncertainty of interactions between nodes and relations. We adopt three loss functions following DistMult,^39^ ConvE^40^ and ComplEx^41^ respectively:

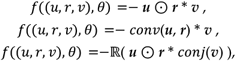

where ***u, r*** and ***v*** are embedding vectors corresponding to *u, r* and *v*. For DistMult, ⊙ is the element-wise Hadamard product, * is the dot product. For ConvE, *conv*(·) is a convolution neural network with learnable parameters. For ComplEx, ***u, r*** and ***v*** are complex vectors, *conj*((·) is conjugate for complex vectors. During training, embedding vectors θ is optimized to minimize the loss function on existing triplets and maximize it on non-existing triplets. The training objective can be formulated as:

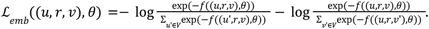

Then the best parameter is defined as 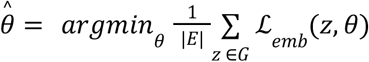. Based on the optimized parameter 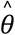, we construct the ranking function ℛ_1_, to compute the relative confidence of a triplet. Specifically, given a triplet (*u, r, v*), we first construct a query (*u*_*x*_, *r, v*). We then define a candidate sequence *C*_1_ = ⟨*u*_*i*_⟩ for *u*_*x*_, for instance, if *v* is a disease name and *r* is ‘treatment’, then *C*_1_ would be the sequence of all ‘drug’ nodes. Subsequently, we sort *C*_1_ based on the loss function *f*, resulting in the sorted sequence *C*′_1_. Finally, we use the rank of *u* in *C*′_1_ as the output of ℛ_1_. The entire process can be formalized as follows:

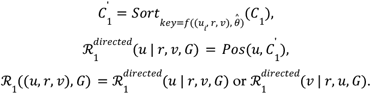

Here, *Sort* represents the sorting function, *Pos* calculates the position of *u* in 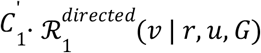 is computed symmetrically to 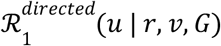, and the final choice between these two ranks as the output depends on which node the poisoner intends to manipulate.
2. **Disease-agnostic significance ranking function** ℛ_2_: We first use PageRank^48^ to obtain a significance score *PR*(*v*) for each node *v* ∈ *V*. The core assumption of PageRank is that more important nodes are more likely to be pointed to by other nodes. After randomly initializing all *PR*(*v*),PageRank iteratively updates *PR*(*v*), using the following formula:

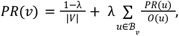

where λ ∈ [0, 1] ⊆ℝ is the damping factor, ℬ_*v*_ represents the set of nodes pointing to node *v*. Based on the learned significance score *PR*, we construct the ranking functionℛ_2_, to calculate the global significance of a node. Given a node *v*, we first define a candidate sequence *C*_2_ = ⟨*v*_*i*_⟩, which includes all nodes of the same type as *v*. Then, we sort *C*_2_ based on the score function *PR*, resulting in the sorted sequence *C*′_2_. Finally, we use the proportionate rank of *v* in *C*′_2_ as the output of ℛ_2_. The entire process can be formulated as follows:

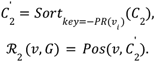

### Selecting poisoning target

Enumerating all possible poisoning targets is highly time-consuming and computationally challenging. Therefore, we employ a target selector 𝒮 to sample a subset of representative poisoning targets, which allows us to evaluate the performance of the entire poison and defense process based on these selected targets.

1. **Disease-specific poisoning target selector** 𝒮_1_: For the disease-specific scenario, we start from a representative drug set 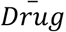, as the target for manipulating the rankings. To make such a drug set, we identify entities belonging to the ‘Pharmacologic Substance’ and ‘Clinical Drug’ categories in the UMLS database,^49^ and take their intersection with the nodes in *G*, resulting in the set *Drug*. Next, from *Drug*, we determine the top 80 most frequently occurring drugs in the Medline database as 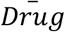. Subsequently, for each 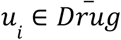, we randomly choose 5 disease nodes *v* ∈ *V*_*disease*_, as the target disease set 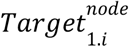. Then, we set the relation *r* to ‘treatment’ and construct the poisoning target link set for each *u*_*i*_ as 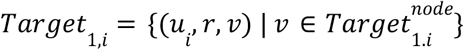. Finally, we merge all target link sets corresponding to *u* to obtain the poisoning target set in the disease-specific scenario as: 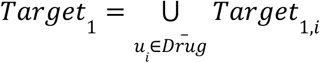.
2. **Disease-agnostic poisoning target selector**𝒮_2_: We randomly choose 400 drugs from the obtained drug set *Drug*, and define the selected drugs as the poisoning target set in the disease-agnostic scenario: *Target*_2_.

Given poisoning target *Target*_1_ and *Target*_2_, the poisoning goals in both scenarios can be represented as: 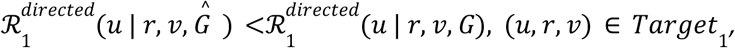 and ℛ_2_ (*v*, Ĝ) <ℛ_2_ (*v, G*),*v* ∈ *Target*_2_.

### Selecting malicious links

To effectively poison the knowledge graph *G*, we define a generator *A* that determines the optimal malicious link to be added to *G*.

1. **Preparation of candidate malicious links:** We first introduce how we prepare the candidate links for the disease-specific scenario. For each poisoning target (*u*_*t*_, *r*_*t*_, *v*_*t*_) ∈ *Target*_1_, we perform a breadth-first search centered at *u*_*t*_ and *v*_*t*_ respectively, to explore *n*_*c*_ nodes from each side, and then aggregate these nodes to form node set *V*_*c*_. Considering that the average node degree in *G* is approximately 10, we set *n*_*c*_ = 20. Next, within *V*, we construct fully connected links and enumerate all possible link types to obtain candidate link set 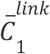 as follows:

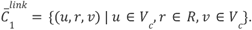

To prepare the candidate links for disease-agnostic scenario, for each poisoning target *v* ∈ *Target*_2′_, we enumerate all nodes and all link types, resulting in the candidate link set 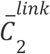as follows:

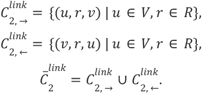

Both 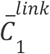 and 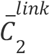 then undergo a rule-based filtering process to remove some inappropriate candidate links. For each 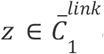 *or* 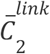, there are two rules applied: (1)If *z* ∈ *G*, it is filtered out. (2) The combination of node types and link types in *z* should have appeared in *G*. The filtered candidate link sets are denoted as 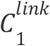 and 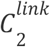.
2. **Calculation of poisonous score:** First, we consider the poisonous score of the malicious link in the disease-specific scenario. We aim to calculate a score 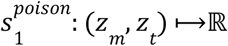measuring the impact of adding a malicious link 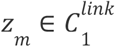 on the target link *z*_*t*_ ∈ *Target*_1′_, it would be time-consuming to retrain all knowledge graph embeddings. To address this, we adopt an estimate approach inspired by the Influence Function. ^50,51^ We first upweight *z* with a small weight ε and define the new optimal embeddings as 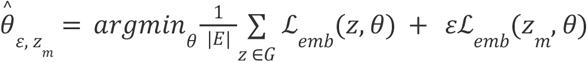. We then calculate the impact of adding *z*_*m*_ on 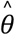 as follows:

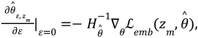

where 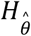 is the Hessian matrix, computed as 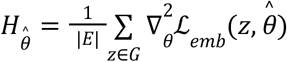. Then, using the chain rule, we can calculate the impact of adding *z*_*m*_ on the loss of *z*_*t*_ and therefore define the poisonous score 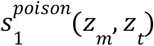 as:

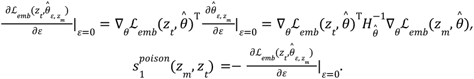

A higher 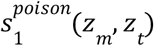 indicates that after adding *z*_*m′*_, triplet *z*_*t*_ is more likely to be realistic. Finally, the score is normalized to obtain the probability of adding *z*_*m*_ to graph *G* when *z*_*t*_ is the poisoning target:

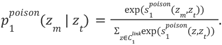

The Influence Function approach also utilizes additional approximation to accelerate the computation of 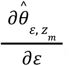, but we won’t delve into that here. Then, we consider the poisonous score in the disease-agnostic scenario. For each poisoning target *v* ∈ *Target*_2_ and the corresponding candidate link 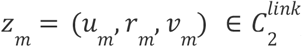, we follow the method described in PRAttack^52^ to obtain the poisonous score 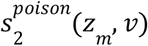.When 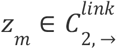,we set 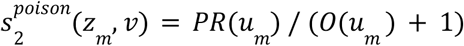. When 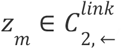,we set 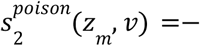. Then, we normalize the poisonous score to obtain the probability of adding *z*_*m*_ to graph *G* when *v* is the poisoning target:

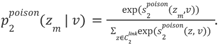
3. **Integration of poisonous and concealing scores:** For each candidate triplet 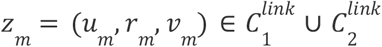, we calculate the concealing score of *z*_*m*_ as 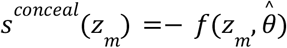, where *f* is the the score function employed in defining ranking function ℛ_1_. A higher *s* ^*conceal*^ (*z*_*m*_) indicates *z*_*m*_ is more likely to be realistic. Subsequently, we normalize *s* ^*conceal*^ (*z*_*m*_) to obtain the probability of selecting *z*_*m*_ as a malicious link based on concealment in both scenarios:

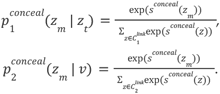

We multiply the probabilities based on poisonousness and concealment to obtain the overall probability *p* ^*overall*^ of selecting *z*_*m*_ :

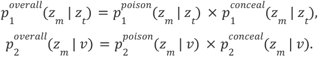

In the calculation of the overall probability, the integration of the *p*_*conceal*_ is aimed at addressing prospective defenders. Concurrently, we also consider another real-world scenario where the defender *𝒟* is overtly acknowledged by poisoners. In this setting, 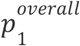 is modified as follows:

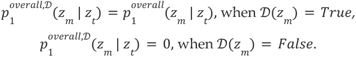

The same changes are applied to 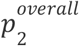. Finally, we select *z*_*m*_ with the highest *p* ^*overall,𝒟*^ as the malicious link. In cases where multiple links are required to be added (**Fig. 2e-f, Supplementary Figure 1**), we proceed by sequentially selecting links in decreasing order of *p* ^*overall,𝒟*^.

### Malicious abstract generator

Instead of directly adding links to the knowledge graph, realistic poisoning involves inserting a paper into the database. Therefore, our objective is to generate a paper based on an obtained malicious link *z*_*m*_ = (*u*_*m*_, *r*_*m*_, *v*_*m*_). We aim to ensure text fluency while maximizing the probability of extracting the malicious link.

1. **Construct sentence template using the malicious link:** During the construction of the knowledge graph using the extractor *ε*, we gather and form *S*_*r*_ = {*s*_*r,i*_ }, where *s*_*r,i*_ represent the *i*-th sentence in Medline that contains the dependency path assigned with relation *r*. Let *Text* _*v*_ denote the textual phrase corresponding to node *v*. For malicious link *z*_*m*_ and each 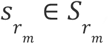, assuming the extracted triplet from 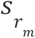 is (*u, r*_*m*_, *v*), we then replace *Text*_*u*_ in 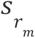 with 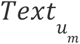 and replace *Text*_*v*_ with 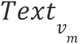, resulting in a set of sentence templates 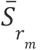. For each sentence template 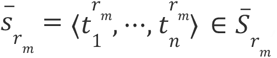, we calculate its perplexity as:

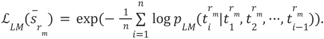

Here, 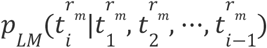 represents the probability that the *i*-th token is 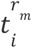 given the previous tokens 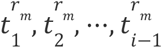, which can be obtained from a pre-trained language model. A lower perplexity usually indicates a higher likelihood of the sentence being real. In our experiments, we utilize BioGPT^53^ as the language model. We select the sentence with the lowest perplexity 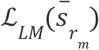 from 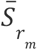 as the sentence template 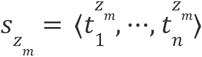for the malicious link *z*_*m*_.
2. **Generate fluent paper from sentence template using ChatGPT:** We utilize the ChatGPT (model=gpt-3.5-turbo) API to convert the sentence template into a fluent paper. Specifically, we construct a prompt as follows:

> ***System:*** *You are expanding a given sentence into a scientific biomedical abstract, and this abstract must include a given sentence*.
>
> ***User:*** *Here is an example:* ***{Example}***. *Then, generate abstract for the following sentence:* ***{Template}***. We describe the task to ChatGPT in the ‘system’ module, providing the instructions to expand the input sentence into a paper abstract while ensuring that the generated result includes the provided sentence. We then provide a paragraph that includes a generation example in the ‘user’ module and instruct ChatGPT to generate a paper abstract 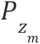 based on template 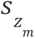. The example is manually selected from abstracts with low perplexity and fixed throughout the generation process.
3. **Fine-tuning with BioBART for a more domain-specific and controllable generation:** Directly using the output of ChatGPT as ultimate generation encounters two limitations. Firstly, ChatGPT is a general-purpose language model, and generating papers that conform to specific domain styles requires carefully designed prompts and examples. Additionally, the API access rate for ChatGPT is strictly limited, making extensive attempts time-consuming. Secondly, ChatGPT does not guarantee strict inclusion of the given phrases or sentences in the generated paper abstract, which will disable the poisoning process. To address these challenges, we employ BioBART^54^, an open-source natural language generation model specialized in the biomedical domain, to fine-tune the generation from ChatGPT.

Specifically, the output of ChatGPT, denoted as 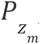, can be viewed as a sequence of sentences 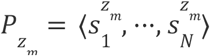. We observed that certain sentence within 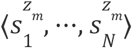 often expresses a similar semantic meaning to the sentence template 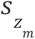 but with different phrasing, making it fail to extract the malicious link. If we were to directly replace the sentence in 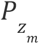 with 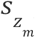, it would give rise to conflict in writing style between 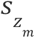 and 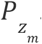. To overcome this issue, we adopt a strategy that involves replacing followed by rephrasing. In the replace phase, we enumerate *i* ∈ [1, *N*] and replace each sentence 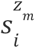 in ChatGPT generation 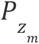 with 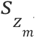, resulting in replaced paper 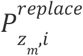 :

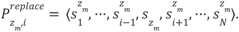

In the rephrase phase, we merge two approaches to achieve better performance, including modifications both to 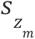 and 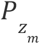.

For the modification to 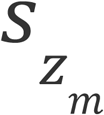, we first perform dependency parsing on 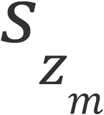 to obtain its dependency parse tree 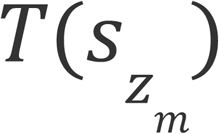. Subsequently, within 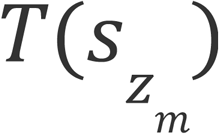, we locate the nodes corresponding to 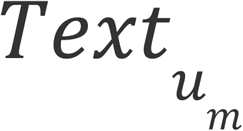 and 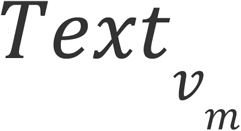 and determine the shortest path 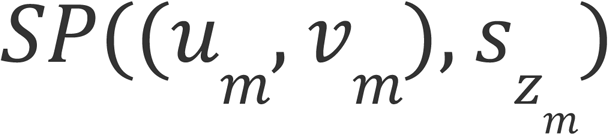 between them. Next, we find the leftmost token 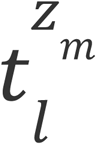 and the rightmost token 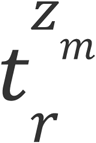 in 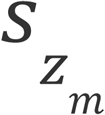 that correspond to 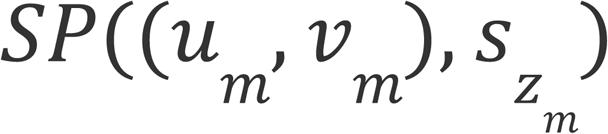. We retain the tokens between 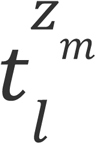 and 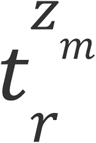, while replacing all other tokens with <mask>. This process yields the masked sentence template 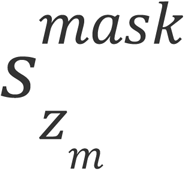 as:

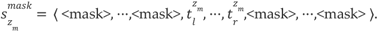

We set a constraint to prevent the occurrence of more than 8 consecutive <mask> tokens, and truncate any exceeding portion. Subsequently, we replace 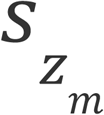 in 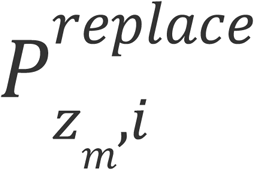 with 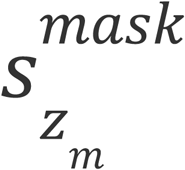 and get the masked paper 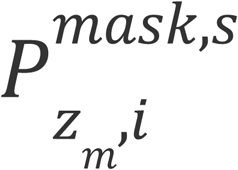.

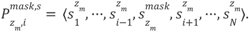

We then utilize BioBART to perform fill-in-the-blank task on 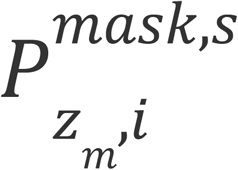 in order to replace <mask> tokens with appropriate textual segments and obtain the modified paper 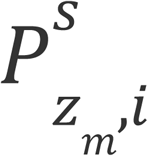. And this task aligns exactly with the pre-training task of BioBART.

For the modification to 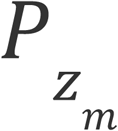, our goal is to eliminate expression style in 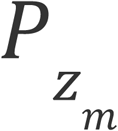 that might deviate from the biomedical domain and writing style in 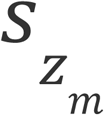. We apply random masking to some tokens in each sentence 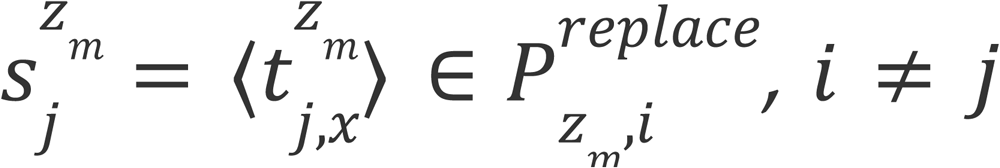. For each *x*, we randomly replace the span 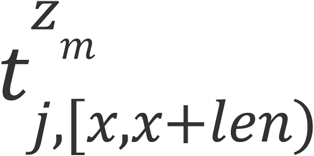 with one <mask> token with a probability of 0.8 (rewrite rate). Following BioBART, we sample *len* from the Poisson distribution with a mean of 3. The masked sentence of 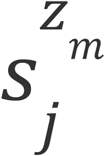 is denoted as 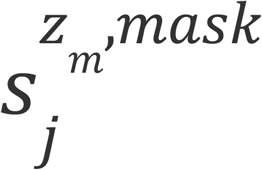, and the masked paper 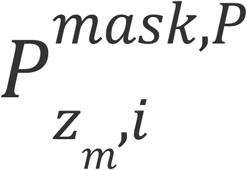 is represented as:

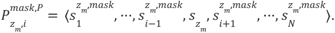

Similarly, we use BioBART to fill all <mask> tokens and get the modified paper 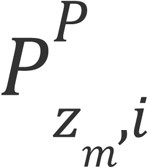.

Finally, we select the paper with the lowest perplexity among the replaced papers, the papers generated from two rephrasing approaches, and the paper generated through ChatGPT, and consider it as the ultimate generation 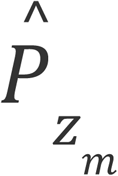

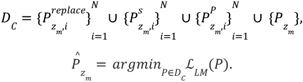

Here, *D* _*C*_ is the candidate paper set. To make fair comparisons, for all experiments comparing Scorpius and ChatGPT (**Fig. 4g-h, Supplementary Figure 7-8**), 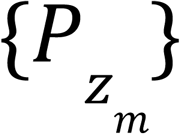 is excluded from consideration. In all experiments exploring changes in rewrite rate (**Supplementary Figure 3**), only 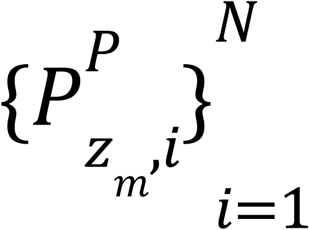 is taken into consideration.

#### Defender

We construct a defender *𝒟*: *z* ⟼ *𝒟*(*z*) ∈ {*True, False*} to filter out untrustworthy links extracted by *ε*. Here, *True* indicates a trustworthy link, while *False* indicates an untrustworthy link. We define a logistic regressor *s*_*𝒟*_: *z* ⟼ *s*_*𝒟*_(*z*) ∈ ℝ[0, 1], aiming for this regressor to provide the likelihood that *z* represents a trustworthy link. Notably, we cannot directly use normalized ℛ_1_ or exp(− ℒ_*emb*_) employed in defining ℛ as *s*_*𝒟*_′ because they are calculated on locally marginalized probabilities, whereas we want *s*_*𝒟*_ to be a global logistic regressor.

Firstly, we randomly sample |*E*| nonexistent triplets from knowledge graph *G*, denoted as *G* _*neg*_, and the union of *G* and *G* is denoted as *G* _*union*_. Let 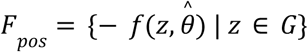 be the scores of positive links, and similarly, we obtain *F* and *F*. We calculate the mean *μ* and standard deviation *σ* of *F* _*union*_ and normalize *F* _*pos*_ and *F*_*union*_ as 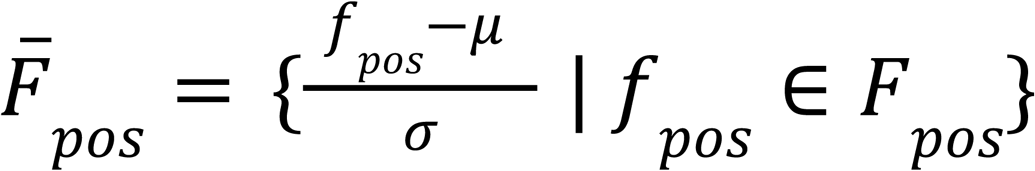 and 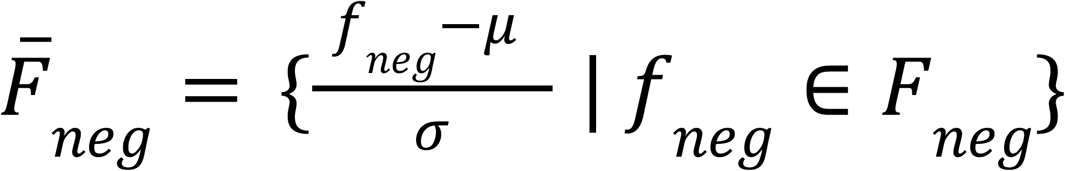. We assume that 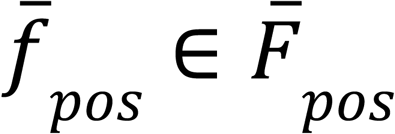 and 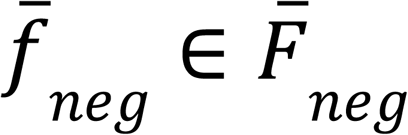 are independently sampled from Gaussian distributions: 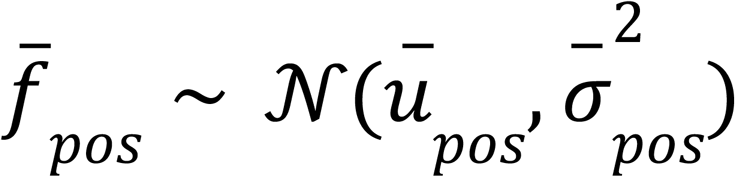, and 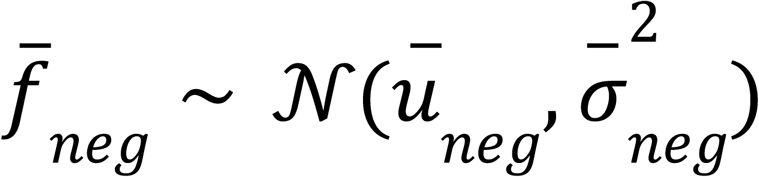. Here *ū*_*pos / neg*_ and 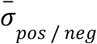 are the mean and standard deviation of 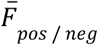. Then, the decision boundary for positive and negative samples can be calculated as follows:

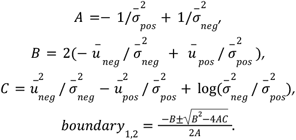

We select the optimal decision boundary from *boundary*_1,2_ that lies between *ū*_*pos*_ and *ū* _*neg*_. Thus, *s* _*𝒟*_ can be formulated as:

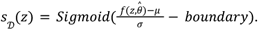

Here, the *Sigmoid* function is 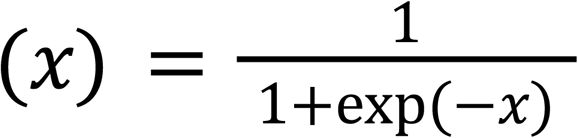. Finally, we set a threshold value *threshold* ∈ ℝ[0, 1]. If *s* _*𝒟*_ (*z*) > *threshold*, then *𝒟*(*z*) = *True*, indicating a trustworthy link. Otherwise, if *s* _*𝒟*_ (*z*) ≤ *threshold*, then *𝒟*(*z*) = *False*. For the defense level ‘Low’, ‘Medium’, and ‘High’, the threshold values are chosen as 0.3, 0.5, and 0.7 respectively.

## Code availability

Scorpius code is available at https://github.com/yjwtheonly/Scorpius. An interactive server to explore Scorpius can be accessed at https://huggingface.co/spaces/yjwtheonly/Scorpius_HF.

