## Supplementary figures and images for "Poisoning scientific knowledge using large language models"

### Supplementary Figure 1

##### Supplementary Figure 1

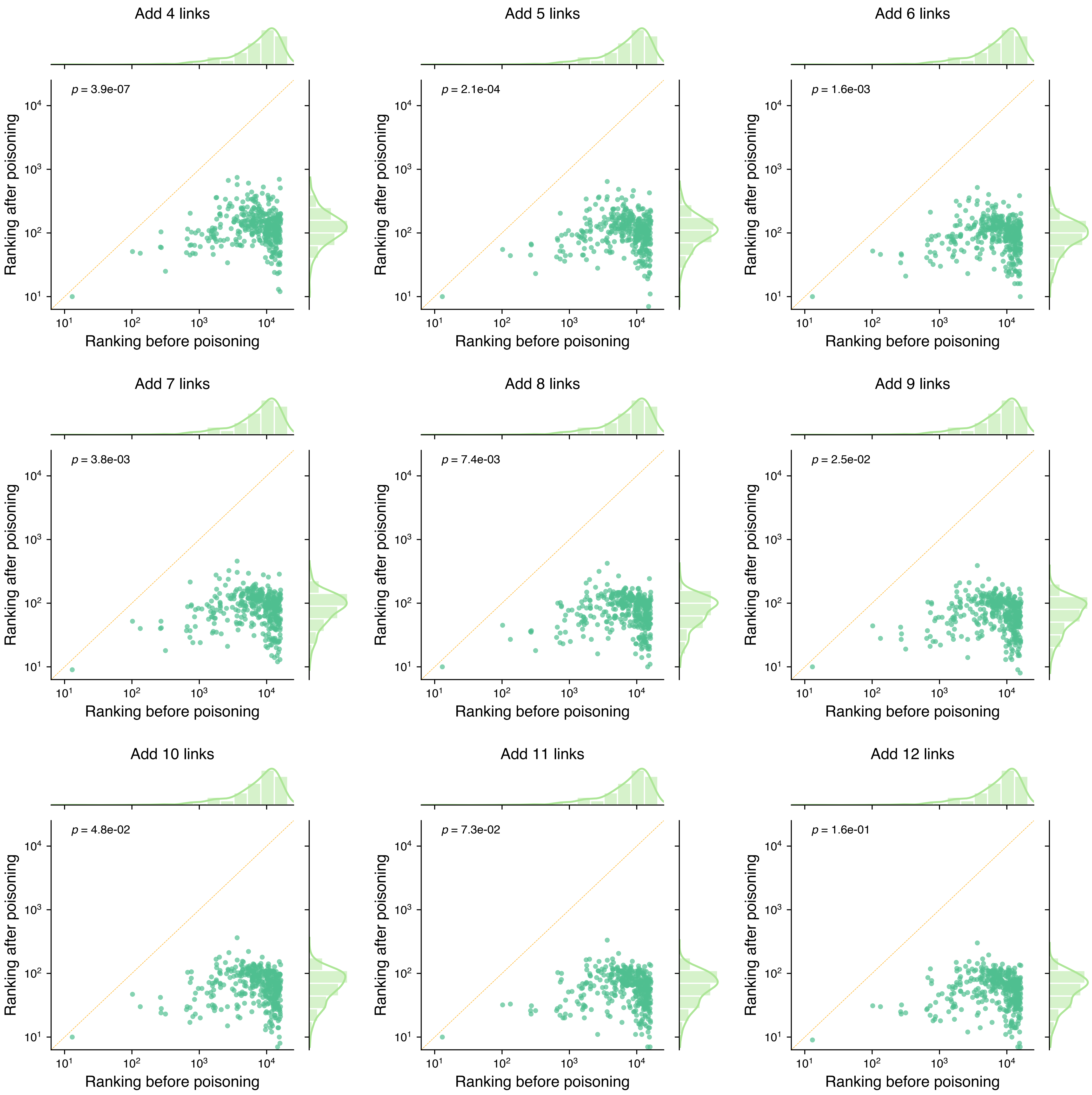

### Supplementary Figure 3

#### Supplementary Figure 3

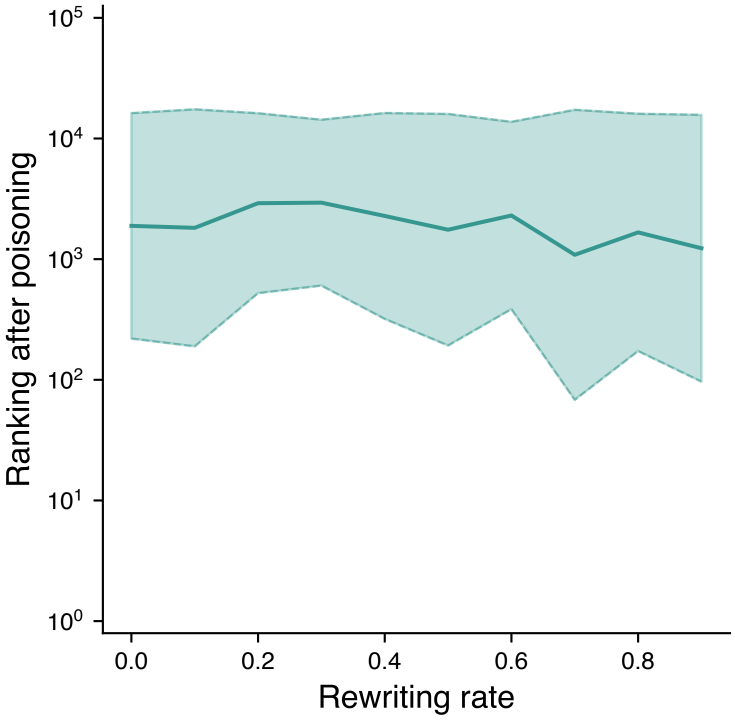

### Supplementary Figure 4

#### Supplementary Figure 4

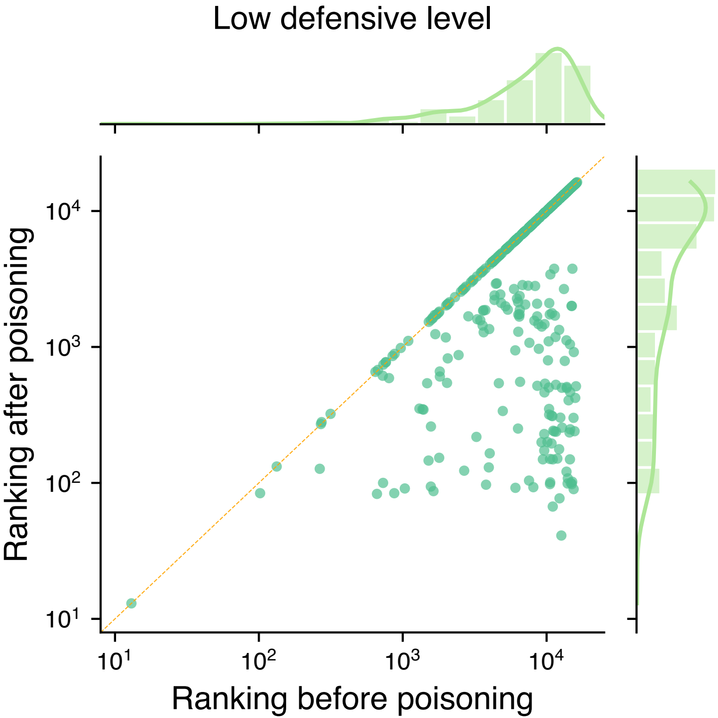

### Supplementary Figure 5

#### Supplementary Figure 5

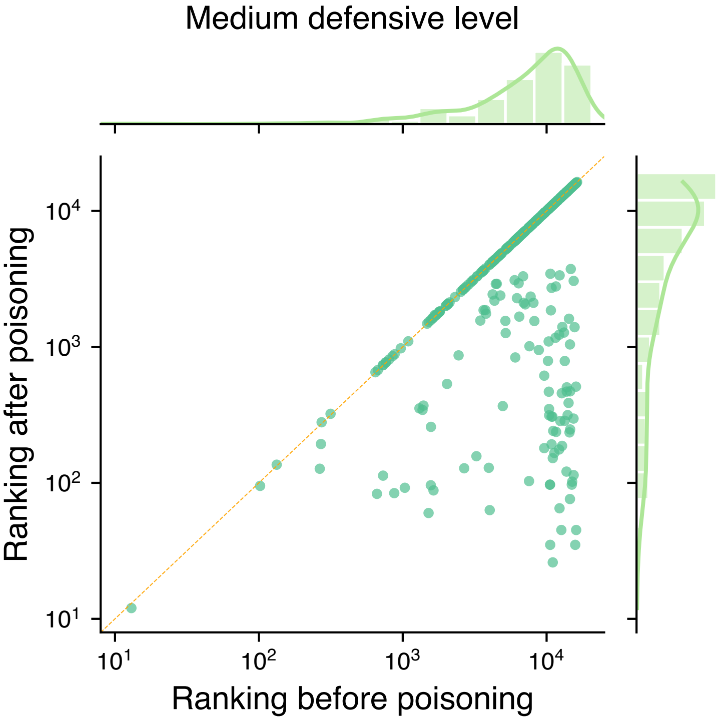

### Supplementary Figure 6

#### Supplementary Figure 6

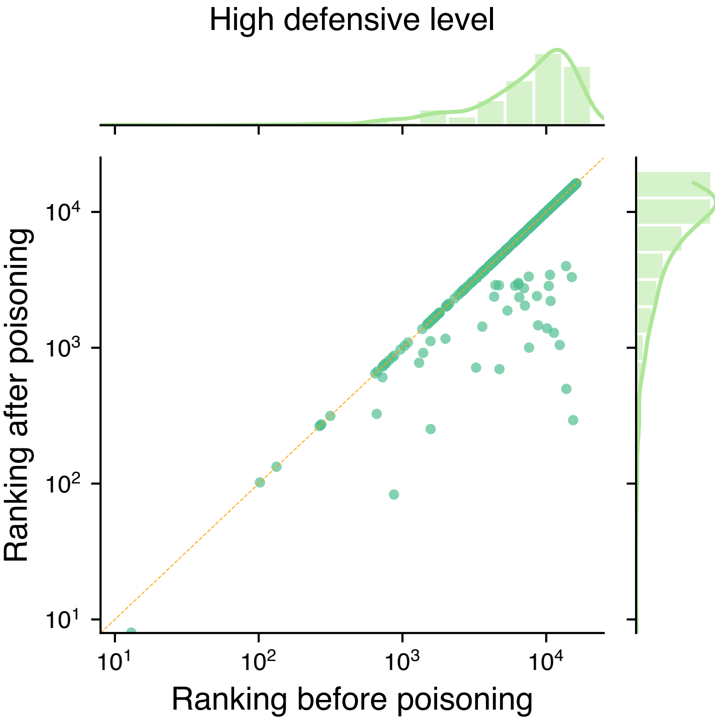

### Supplementary Figure 7

#### Supplementary Figure 7

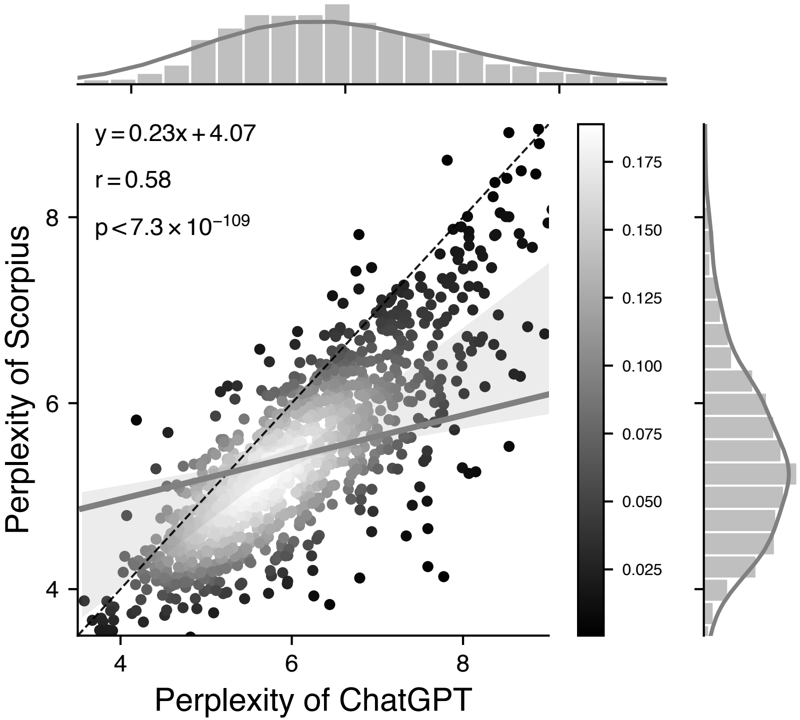

### Supplementary Figure 8

#### Supplementary Figure 8

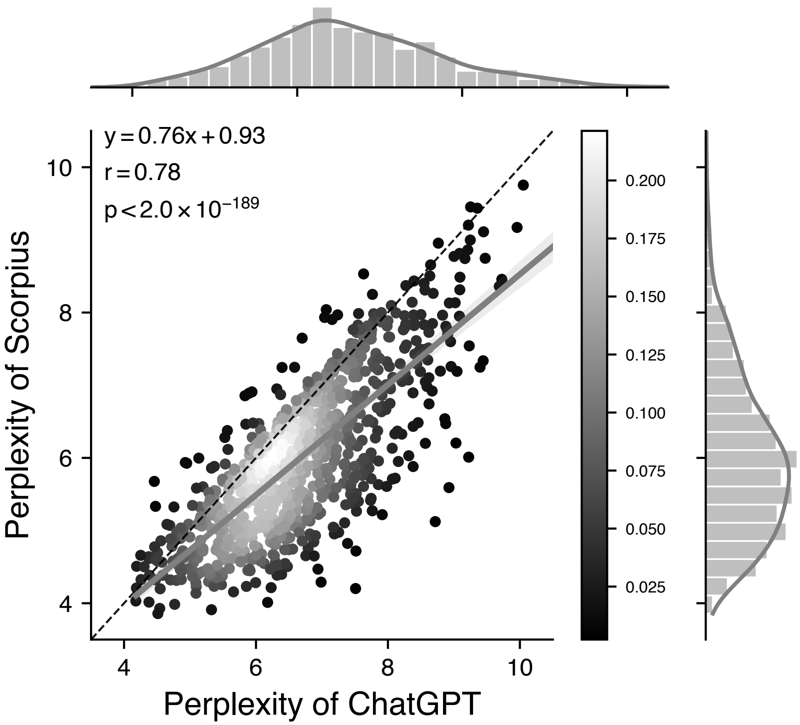
