## Supplementary Figure 2 for "Poisoning scientific knowledge using large language models"

In summary, our results demonstrate that MG132 induces the pro-apoptotic protein cotransin by a p53-independent mechanism that leads to caspase-dependent apopt apopt.

20% replacement rate

In summary, our results show that MG132 induces the degradation protein cotransin- a p53-independent mechanism that leads to caspase-dependent apopt apopt.

40% replacement rate

In the, our results show in MG132 induces the degradationoph cotransin- a p53-independent, that is c caspase-dependent apopt apopt.

60% replacement rate

Although TMEM-218 and MN have been reported in two the same patient and both have have induced by drug exposures, this is the first report of MN in a patient with TMEM-218 likely induced by levamisole.

20% replacement rate

Although TMEM-218 and MN have been reported reported two the same patient and both same two induced by drug lev, this is to first report of a in a patient patient TMEM-218 likely induced by levamisole.

40% replacement rate

Although TMEM-218 and MN- been shown first first the same compound struct two same same induced by drug induced same this is is first report of the in a single the TMEM-218 induced MN MN levamisole induced.

60% replacement rate

We also studied the effect of MAGEB16 on podocyte nephrin expression, reactive oxygen oxygen ( ROS ) generation ( ( DCFDA loading ( by fluorometric analysis ), proliferation, and apopt ( morphologic assays ).

20% replacement rate

We also studied the effect of MAGEB16 on podocyte nephrin expression, reactive oxygen oxygen ( R ) generation ( ( DCFDA loading ( by fluorometric analysis ), proliferation, and the morph morphologic changes,.

40% replacement rate

We also studied the effect of MAGEB16, podocyte nephrin (, pod pod pod ( ) ) and,, DCFDA loading was by flow analysis of the proliferation, and apopt apopt apopt apopt apopt.

60% replacement rate
